# Assembly of the algal CO_2_-fixing organelle, the pyrenoid, is guided by a Rubisco-binding motif

**DOI:** 10.1101/2020.08.16.252858

**Authors:** Moritz T. Meyer, Alan K. Itakura, Weronika Patena, Lianyong Wang, Shan He, Tom Emrich-Mills, Chun S. Lau, Gary Yates, Luke C. M. Mackinder, Martin C. Jonikas

**Author notes:** Department of Chemical and Systems Biology, Stanford University, Stanford, CA 94305, USA.

## Abstract

Approximately one-third of the Earth’s photosynthetic CO_2_ assimilation occurs in a pyrenoid, an organelle containing the CO_2_-fixing enzyme Rubisco. How constituent proteins are recruited to the pyrenoid, and how the organelle’s sub-compartments - membrane tubules, a surrounding phase-separated Rubisco matrix, and a peripheral starch sheath - are held together is unknown. Using the model alga *Chlamydomonas reinhardtii*, we discovered that pyrenoid proteins share a sequence motif. We show that the motif is sufficient to target proteins to the pyrenoid and that the motif binds to Rubisco, suggesting a mechanism for targeting. The presence of the Rubisco-binding motif on proteins that localize to the tubules and on proteins that localize to the matrix-starch sheath interface suggests that the motif holds the pyrenoid’s three sub-compartments together. Our findings advance our understanding of pyrenoid biogenesis and illustrate how a single protein motif can underlie the architecture of a complex multi-layered phase-separated organelle.

**One Sentence Summary:** A ubiquitous Rubisco-binding motif targets proteins to the pyrenoid and holds together the pyrenoid’s three sub-compartments.

---

CO_2_ is the source of carbon for nearly the entire biosphere (*1, 2*), but its availability is limited in many environments. To overcome this limitation, many photosynthetic organisms use energy to locally concentrate CO_2_ around the CO_2_-assimilating enzyme Rubisco (*3–6*). In eukaryotic algae, which perform a major fraction of primary production in the oceans (*7*), concentrated CO_2_ is supplied to Rubisco inside a micro-compartment, the pyrenoid (*8*). The pyrenoid consists of a spheroidal protein matrix containing Rubisco, which in nearly all algae is traversed by membranous tubules through which CO_2_ is delivered (*9, 10*). In green algae, the Rubisco matrix is additionally surrounded by a starch sheath (*11*) that limits CO_2_ leakage out of the pyrenoid (*12*) (Fig. 1A).

**Fig. 1.**
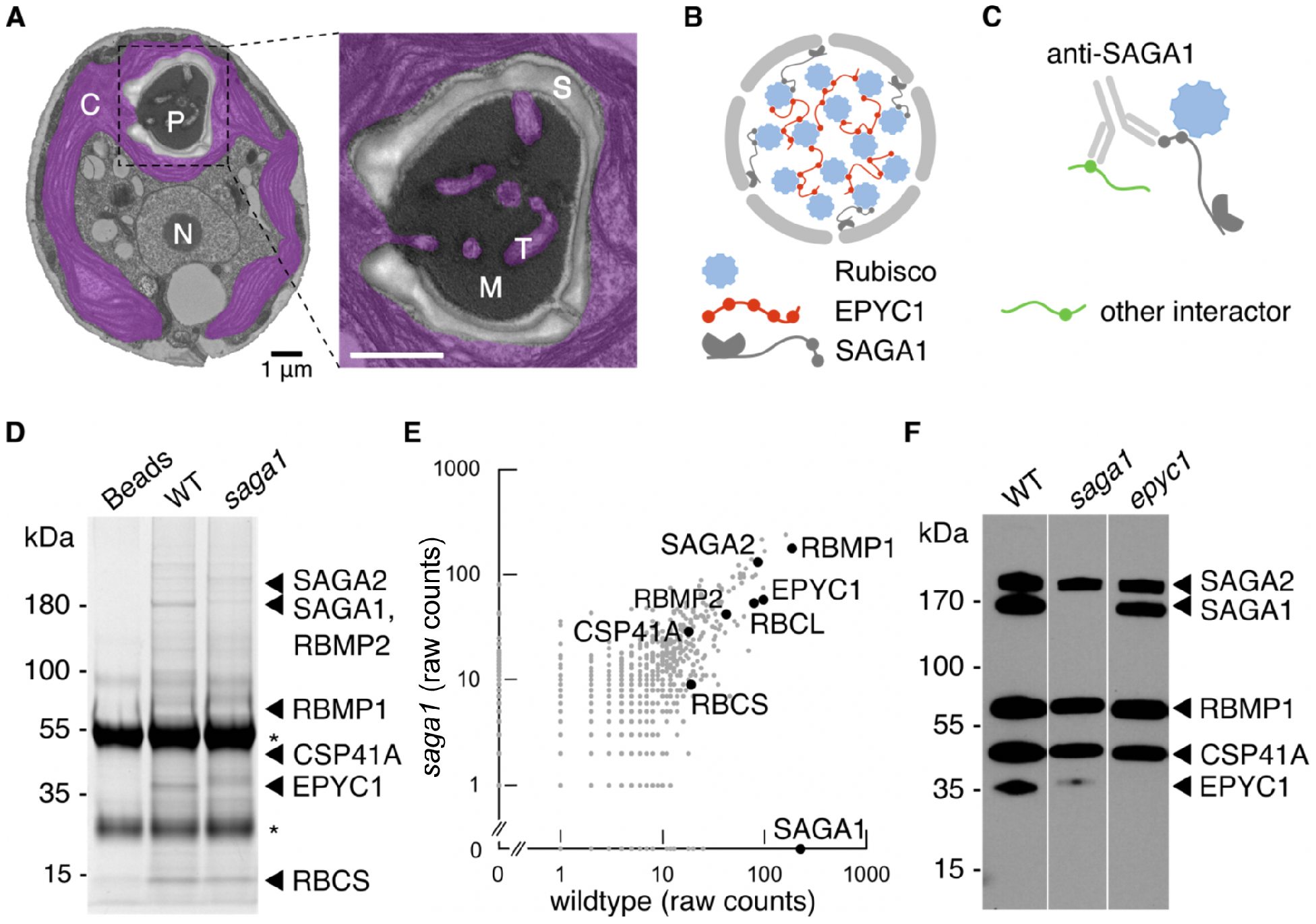
A polyclonal antibody raised against the pyrenoid protein SAGA1 interacts with at least five other pyrenoid proteins. (**A**) Electron micrograph of a median plane section through an air acclimated Chlamydomonas cell. N: nucleus; C: chloroplast; P: pyrenoid; M: Rubisco matrix; T: tubules; S: starch sheath. Scale bar, 1 μm. (**B**) Two proteins, the Rubisco linker EPYC1 and the starch sheath-binding protein SAGA1, have been previously characterized and localized to the pyrenoid. (**C**) An anti-SAGA1 antibody was incubated with cell lysate in an effort to coimmunoprecipitate proteins that bind to SAGA1. (**D**) Coomassie stained SDS-PAGE of proteins immunoprecipitated by the anti-SAGA1 antibody from wildtype (WT) and *saga1* mutant lysates. Immunoprecipitated proteins were eluted from anti-SAGA1 antibodies on beads by boiling; beads not incubated with lysate were also boiled for reference. (*): heavy and light immunoglobulin chains. (**E**) Proteins immunoprecipitated by the SAGA1 antibody from wildtype and *saga1* were identified by mass spectrometry. Raw spectral counts are plotted on a log scale. (**F**) Anti-SAGA1 western blot on denatured protein extracted from wildtype, *saga1* and *epyc1*.

In the model green alga *Chlamydomonas reinhardtii* (Chlamydomonas hereafter), two proteins are known to play central roles in pyrenoid assembly, EPYC1 (*13*) and SAGA1 (*14*) (Fig. 1B). EPYC1 is a ~35 kDa intrinsically disordered protein that phase-separates with Rubisco to form the pyrenoid matrix (*13, 15, 16*). SAGA1 is a ~180 kDa protein with a starch binding domain that localizes to the periphery of the matrix and is required for normal pyrenoid morphology, although the underlying molecular mechanism is unknown (*14*). Here, we present how further characterization of SAGA1 led us to discover that pyrenoid-localized proteins share a common Rubisco-binding protein sequence motif, providing insights into protein targeting to the pyrenoid and assembly of the pyrenoid’s three sub-compartments.

In an effort to immunoprecipitate SAGA1 interacting proteins, we incubated a polyclonal SAGA1 antibody with clarified cell lysates from wildtype cells (Fig. 1C; Fig. S1). In addition to SAGA1, the antibody precipitated multiple proteins found in the pyrenoid proteome (*17*), suggesting at first that those proteins may interact with SAGA1. The precipitated proteins included EPYC1, SAGA2, RBMP1, RBMP2 and CSP41A (Fig. 1D and 1E; Table S1). The protein we named SAGA2 (Cre09.g394621) is 30% identical to SAGA1 (Cre11.g467712) and, like SAGA1, has a predicted starch-binding domain (Fig. S2A to S2C). The proteins we named Rubisco-Binding Membrane Proteins RBMP1 (Cre06.g261750) and RBMP2 (Cre09.g416850) have predicted transmembrane domains and were previously found to bind to Rubisco (*18*) (Fig. S2D to S2G). CSP41A (Cre10.g440050) is a predicted chloroplast epimerase (*19*) (Fig. S2H).

Surprisingly, we found that the precipitation of these pyrenoid proteome proteins was not mediated by SAGA1, but rather that the proteins were directly bound by the SAGA1 antibody. Three lines of evidence supported this conclusion. Firstly, the same proteins were immunoprecipitated from a *saga1* mutant lysate (Fig. 1D and 1E; Table S1). Secondly, the predicted molecular weights of EPYC1, SAGA2, RBMP1, RBMP2, and CSP41A showed remarkable agreement with the multiple polypeptides recognized by the same SAGA1 antibody in immunoblots (*14*) (compare Fig. 1D and 1F). Thirdly, the ~35 kDa band was absent in anti-SAGA1 immunoblots of *epyc1* mutant cell extracts, strongly suggesting that EPYC1 was directly recognized by the SAGA1 antibody (Fig. 1F). We therefore hypothesized that our polyclonal SAGA1 antibody recognized similar sequences on all six pyrenoid proteins, raising questions about the nature and function of this sequence.

To identify potential sequences in the proteins that our antibody could bind, we searched their sequences for similarity to the nineteen C-terminal amino acids of SAGA1, against which our antibody had been raised (*14*). Sequence alignment revealed that all six proteins contain a common motif, with sequence [D/N]W[R/K]XX[L/I/V/A], present at their exact C-termini (Fig. 2A and 2B).

**Fig. 2.**
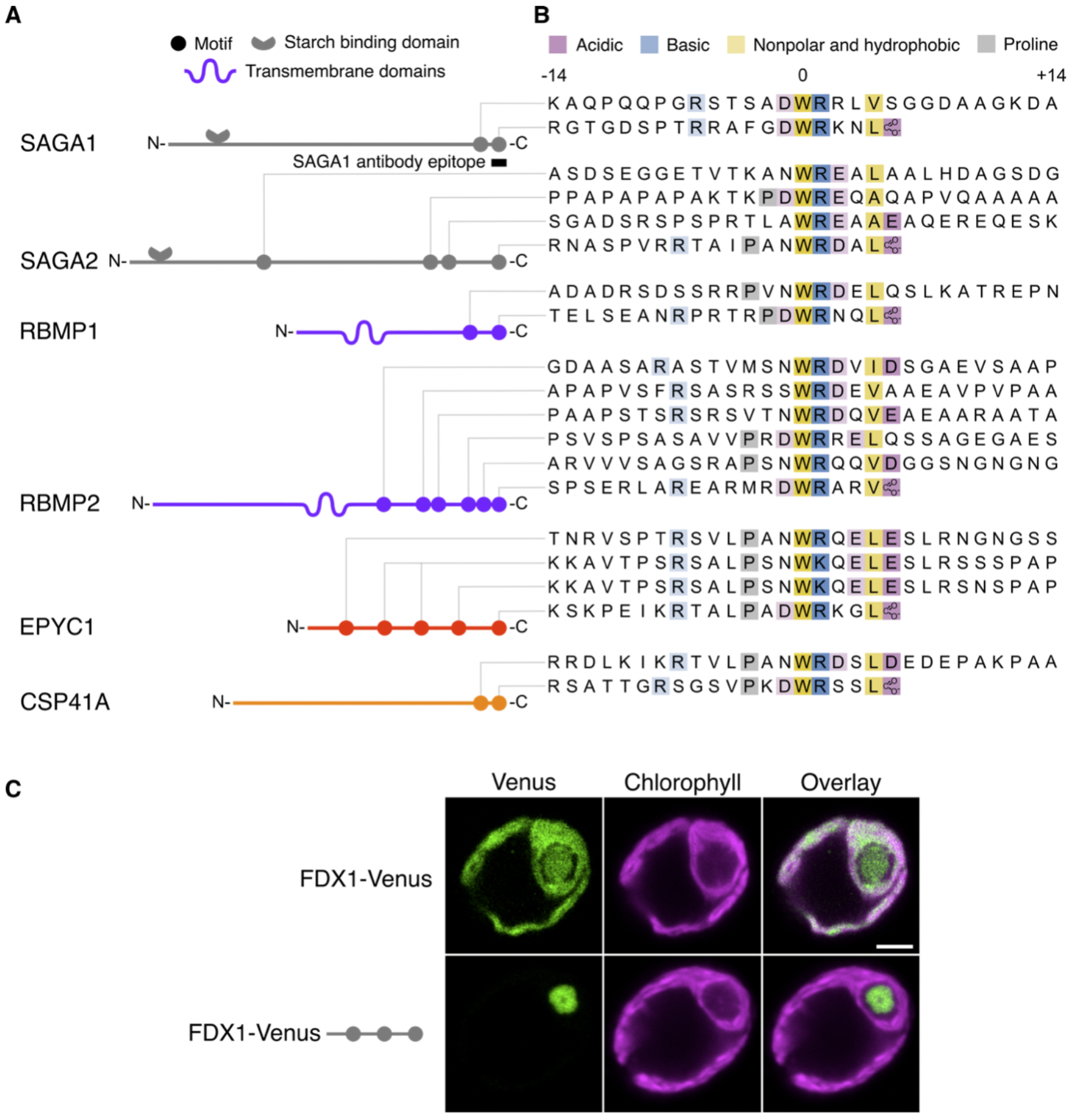
A motif found on pyrenoid proteins is sufficient for targeting proteins to the pyrenoid. (**A**) The location of motifs along the primary sequence of each protein is shown (not to scale). (**B**) Sequence alignment of protein regions containing the pyrenoid motif. Motif residues are colored by physicochemical properties. Intensity of coloring is proportional to frequency at a given amino acid position. Peptides with the sequences shown in **B** were synthesized and their binding to Rubisco was measured by surface plasmon resonance and peptide tiling array (see Fig. 3). (**C**) The localization of Venus-tagged ferredoxin 1 protein (FDX1) without and with the C-terminal addition of three copies of the C-terminal SAGA2 motif was determined by transforming the corresponding constructs into wildtype Chlamydomonas. Scale bar, 2μm.

We identified an additional 14 variants of this motif at internal positions across all six proteins (Fig. 2A and 2B). Most internal occurrences of the motif are immediately followed by an aspartic acid (D) or a glutamic acid (E), both of which contain a carboxyl group. This observation suggests that when the motif is found at the C-terminus of the protein, the C-terminal carboxyl group of the protein is functionally important; and when the motif is found internally in the protein, this functionality is provided by the carboxyl group of the D or E side chains that follow the motif.

In summary, all six of the pyrenoid proteins share multiple copies of a common motif, which appears to have been coincidentally recognized by our SAGA1 antibody.

The prevalence of a common motif among pyrenoid-localized proteins led us to hypothesize that this motif mediates targeting of proteins to the pyrenoid. To test this hypothesis, we evaluated the impact of adding the motif to ferredoxin 1 (FDX1, Cre14.g626700), a small soluble protein that natively localizes throughout the chloroplast stroma. To increase the chances of observing the motif’s effect, we chose to add three tandem copies of the motif to FDX1. Strikingly, addition of the motif re-localized fluorescently-tagged FDX1 exclusively to the pyrenoid matrix (Fig. 2C; Fig. S3A and S3B). We obtained a similar result with another chloroplast protein (Fig. S3C and S3E). These results demonstrate that the motif is sufficient to localize a chloroplast protein to the pyrenoid matrix.

Two observations led us to hypothesize that proteins bearing the motif are recruited to the pyrenoid via binding to Rubisco. First, the motif is present in each of the regions that were found to mediate EPYC1’s binding to Rubisco in a parallel study (*20*). Second, the other five motif-containing proteins were also previously found to bind to Rubisco: SAGA1 by yeast two-hybrid (*14*), and SAGA2, RBMP1, RBMP2 and CSP41A by affinity purification-mass spectrometry (*18*).

To determine whether each variant of the motif can bind to Rubisco, we used surface plasmon resonance to measure the binding of synthetic peptides to Rubisco. With this method, 15 out of 20 peptides representing motif variants had a higher affinity to Rubisco than peptides with random sequences. Peptides with C-terminal motifs systematically showed higher affinity to Rubisco than peptides with internal motifs (Fig. 3A). Rubisco bound to all predicted internal motif sites when we incubated purified Rubisco with arrays of peptides tiling across the full-length proteins (Fig. 3B and 3C; Fig. S4A to S4D; C-terminal sites could not be assayed by this method). These results indicate that nearly all instances of the motif bind to Rubisco *in vitro*. Importantly, binding of the proteins to the SAGA1 antibody in the immunoprecipitation experiment (Fig. 1C and 1D) suggests that at least one motif on each protein is accessible for Rubisco binding when the proteins adopt their native folding *in vivo*.

**Fig. 3.**
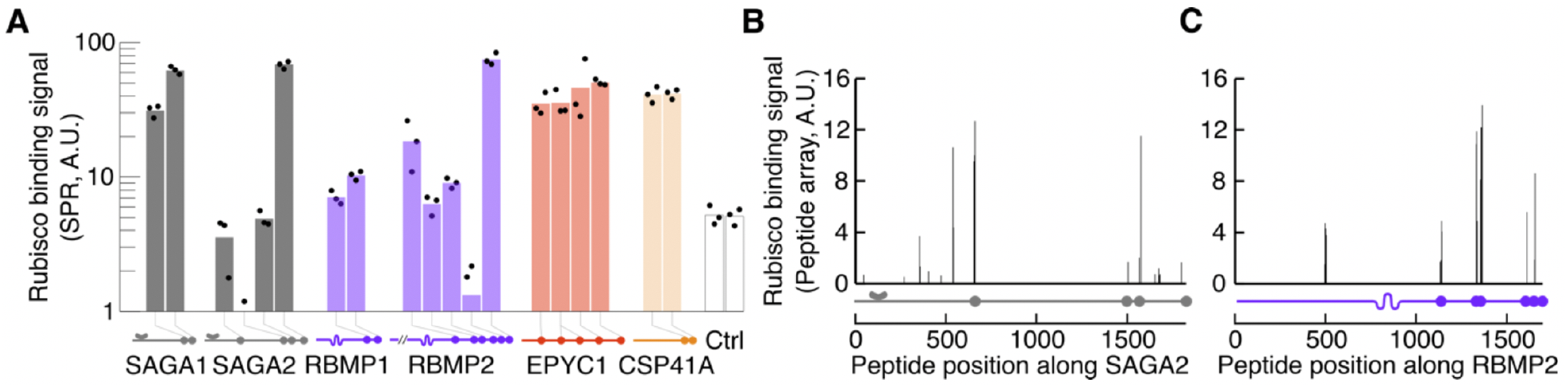
The motif binds to Rubisco. (**A**) Peptides containing motifs from the indicated proteins were synthesized and their binding to Rubisco was measured by surface plasmon resonance (A.U., arbitrary units). Two peptides not containing the motif were included as controls. Internal peptides were 29 amino acids; C-terminal peptides were 19 amino acids; the signal is normalized by peptide molecular weight. The positions of the predicted motifs are indicated below the graph (not to scale). (**B**) Array of 18 amino acid peptides tiling across the sequence of SAGA2 was synthesized and probed with Rubisco. (**C**) Tiling array for RBMP2. Binding signal in (B) and (C) is normalized to a control EPYC1 peptide known to bind to Rubisco (*20*) (one unit of binding). The positions of motifs are indicated to scale below each graph.

Similar motifs are present in proteins other than the ones we studied here. We identified putative motifs in the proteome using a simple scoring scheme (Table S2). Cre10.g430350, a protein with a predicted motif identified by this search, localized to the pyrenoid matrix (Fig. S5A). High motif scores were modestly enriched among pyrenoid proteome proteins relative to other proteins in the chloroplast proteome (Fig. S5B). Not all pyrenoid proteome proteins have the motif; some proteins may be targeted to the pyrenoid via other mechanisms, for example by binding to a motif-containing protein. The motif is present in proteins that are not in the pyrenoid proteome; this could be due to these motifs not being accessible for interaction with Rubisco.

Serendipitously, in a parallel study (*20*) we determined where the motif binds on Rubisco. In that study, as part of an effort to understand how EPYC1 clusters Rubisco to form the pyrenoid matrix, we obtained a cryoelectron microscopy structure of Rubisco bound to a peptide from EPYC1. Remarkably, a Rubisco-binding motif is present at positions N_62_W_63_R_64_Q_65_E_66_L_67_E_68_ on this EPYC1 peptide and plays a central role in the binding interface. One EPYC1 peptide binds to each of the eight Rubisco small subunits of the Rubisco holoenzyme. The motif-containing portion of the peptide adopts an alpha-helix that binds to the Rubisco surface. R_64_ of the peptide, corresponding to the [R/K] of the motif, forms a salt bridge with a glutamic acid of the Rubisco small subunit. Additionally, W_63_ and L_67_ of the peptide, corresponding to the W and [L/I/V/A] of the motif, respectively, contribute to a hydrophobic interaction with three hydrophobic residues of the Rubisco small subunit. The central role of the motif residues in the binding interface strongly suggests that all Rubisco-binding motifs studied here bind to Rubisco using the same mechanism.

Together, our findings suggest a mechanism for targeting proteins to the pyrenoid, involving the presence of a common motif which recruits its protein to the pyrenoid via direct binding interactions with Rubisco. It is possible that the mechanism operates via random diffusion of the motif-bearing protein through the chloroplast, followed by capture of the motif by Rubisco when the protein encounters the pyrenoid matrix.

Beyond providing a mechanism for targeting proteins to the pyrenoid matrix, the motif appears to play a role at the interfaces of the pyrenoid’s three sub-compartments. Whereas Rubisco and EPYC1 are localized in the matrix (Fig. 4A; Fig. S6A and S6B), we observed that some motif-containing proteins localize to pyrenoid regions other than the matrix. Fluorescently tagged RBMP1 and RBMP2 localized to the tubules (Fig. 4A; Fig. S6C and S6D). SAGA1 (*14*) and SAGA2 localized to the interface between the Rubisco matrix and the starch sheath (Fig. 4A; Fig. S6E and S6F).

**Fig. 4.**
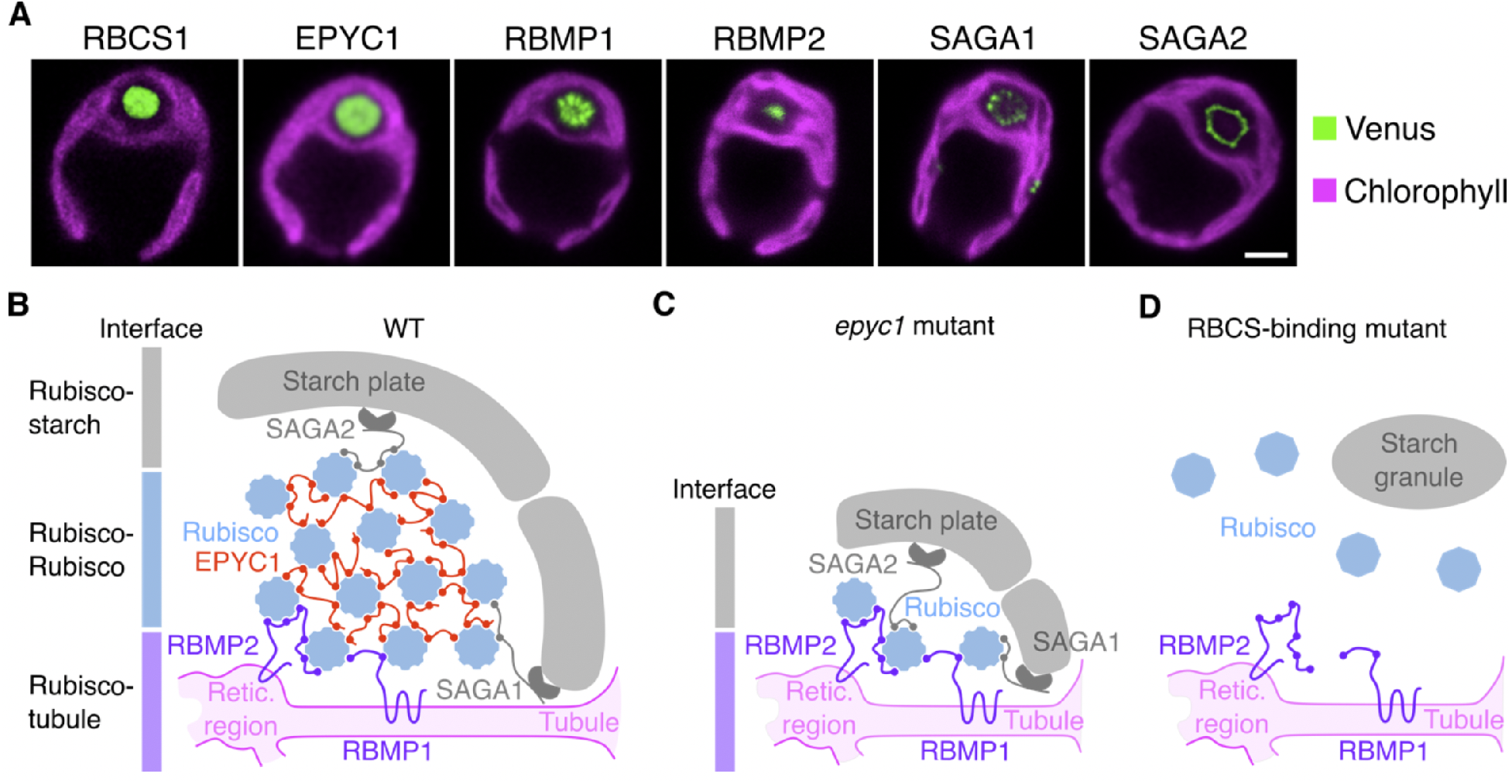
The motif orchestrates the architecture of the pyrenoid’s three sub-compartments. (**A**) Representative confocal images of Venus-tagged proteins that possess the Rubisco-binding motif, and Rubisco small subunit (RBCS1). Chlorophyll autofluorescence delimits the chloroplast. Scale bar, 2 μm. (**B**) Proposed model for how the motif mediates assembly of the pyrenoid’s three sub-compartments in wildtype. The motif on tubule-localized transmembrane proteins RBMP1 and RBMP2 mediates Rubisco binding to the tubules (Retic. region = reticulated region of tubules (*9*)). Multiple copies of the motif on EPYC1 link Rubiscos to form the pyrenoid matrix (*20*). At the periphery of the matrix, the motif on starch-binding proteins SAGA1 and SAGA2 mediates interactions between the matrix and surrounding starch sheath. (**C**) The model in Fig. 4B explains the matrix-less phenotype observed in EPYC1-less mutants. (**D**) The model also explains the absence of matrix and starch plates in mutants where Rubisco’s binding site for the motif has been disrupted (*20*).

These observations together with previous work (*9, 13–17, 20*) are consistent with a model where the Rubisco-binding motif holds together the pyrenoid matrix, the traversing tubules, and the surrounding starch sheath (Fig. 4B). In this model, multiple copies of the motif on EPYC1 mediate cohesion of the matrix by bringing Rubisco holoenzymes together (*20*). Moreover, the presence of the same motif on the pyrenoid tubule-localized transmembrane proteins, RBMP1 and RBMP2, recruits Rubisco to the tubules and favors assembly of the matrix around them. Finally, the presence of the motif on proteins with starch-binding domains, SAGA1 and SAGA2, which localize to the pyrenoid periphery, mediates adherence of the starch sheath to the matrix.

Our microscopy data suggest that RBMP2 is confined to the central reticulated region of the tubules (*9*), whereas RBMP1 appears to localize to the more peripheral tubular regions to the exclusion of the reticulated region. These different localization patterns suggest that RBMP1 and RBMP2 could each promote Rubisco matrix binding to a different part of the tubules. As described previously (*14*), SAGA1 localized to puncta at the periphery of the matrix, likely at the interface between the Rubisco matrix and the starch sheath. We observed that SAGA2 also localized to that interface but appeared to cover the surface of the matrix more homogeneously than SAGA1. Similarly to the RBMPs, the different localization patterns of the SAGAs suggest that they may each interface with different features on the starch sheath.

The model explains several previously puzzling observations. First, in a mutant lacking EPYC1, a pyrenoid-like structure still assembles around the tubules, containing some Rubisco enclosed by a starch sheath, even though the canonical matrix is absent (*13*). The presence of both Rubisco and starch in this structure can be explained by a layer of Rubisco that serves as a bridge between motif-containing proteins RBMP1 and RBMP2 on the tubules and motif-containing proteins SAGA1 and SAGA2 on the surrounding starch sheath (Fig. 4C). Second, point mutations that disrupt the EPYC1 binding site on Rubisco not only eliminate the matrix but additionally disrupt the association of the starch sheath with the pyrenoid (*20*). The additional disruption of the starch sheath can be explained by the same Rubisco binding site being required not only for binding to EPYC1 but also for binding the motif on other proteins that connect the starch and tubules to Rubisco (Fig. 4D).

Our work reveals a ubiquitous Rubisco-binding motif that is both sufficient for targeting proteins to the pyrenoid and also appears to mediate the overall assembly of the pyrenoid’s three sub-compartments. The eight-fold symmetry of the Rubisco holoenzyme allows it to interact simultaneously with multiple binding partners via the motif, making Rubisco a central structural hub of the pyrenoid. The valency and binding strengths of the motif to Rubisco vary among the binding partners (Fig. 2A and Fig. 3), which could play a role in their relative priority of access to Rubisco, as observed for other phase-separated organelles (*21–23*).

The Rubisco-EPYC1 condensate can enhance CO_2_ fixation only if it is anchored around the pyrenoid tubules, which are thought to provide concentrated CO_2_ (*8, 9, 24, 25*). Identification in the present work of two tubule-localized proteins that bind to Rubisco provides a plausible explanation for how the Chlamydomonas pyrenoid forms preferentially around tubules, rather than anywhere else in the chloroplast. Considering that the tubules can form independently of the matrix (*26, 27*), whether the motif is required for RBMPs to localize to the tubules remains to be investigated. Our observation that RBMP1 and RBMP2 localize to different sub-domains of the tubules suggests that an additional localization mechanism may be at play, such as a preference for a specific membrane curvature.

Intriguingly, RBMP1 is predicted to be a Ca^2+^-activated anion channel of the bestrophin family (Fig. S2D and S3F), of which several members are thought to supply HCO_3_^-^ to the lumen of the tubules for conversion to CO_2_ (*28*). Whereas the previously described members localized primarily to membranes outside the pyrenoid, RBMP1 localizes exclusively to the tubules themselves, which may allow it to directly feed HCO_3_^-^ to the tubules for conversion to CO_2_.

We hypothesize that the organizational principle described here applies broadly to pyrenoids across the tree of life. Pyrenoids are thought to have evolved independently multiple times through convergent evolution (*29, 30*). Their convergent evolution may well have been facilitated by the possibility of using a common motif and binding site on Rubisco to perform three functions essential to all pyrenoids: clustering of Rubisco into a matrix, targeting of proteins to the matrix, and connecting the matrix to other structures.

Our findings advance the basic understanding of the biogenesis of the pyrenoid and provide a framework for engineering a pyrenoid into crops for improving yields (*8, 31, 32*). More broadly, the system presented here provides a remarkable example of how the architecture of a complex phase-separated organelle can be defined by a simple organizing principle.

## Acknowledgements

We thank Christopher Dupont (JCVI, La Jolla, CA), as well as present members and alumni of the Jonikas laboratory for helpful discussions and feedback on the manuscript; the Princeton University Confocal Microscopy and Biophysics core facility managers, Drs. Gary Laevsky and Venu Gopal Vandavasi for instrumentation support; Ryan Leib and Christopher Adams at the Stanford University Mass Spectrometry facility.

## Funding

The project was funded by National Science Foundation (IOS-1359682 and MCB-1935444), National Institutes of Health (DP2-GM-119137), and Simons Foundation and Howard Hughes Medical Institute (55108535) grants to M.C.J.; and the UK Biotechnology and Biological Sciences Research Council (BB/R001014/1) grant to L.C.M. The content is solely the responsibility of the authors and does not necessarily represent the official view of the National Institutes of Health.

## Author contributions

M.T.M., A.I. and M.C.J. conceived the project. A.I. performed the coimmunoprecipitation experiment and prepared samples for analysis by mass spectrometry. M.T.M. purified Rubisco and performed western blots, surface plasmon resonance, protein relocalization experiments, and confocal microscopy. S.H. and M.T.M. performed the peptide tiling array experiment. T.E.-M., J.L., G.Y., L.W., M.T.M. and L.C.M. produced fluorescently tagged strains. W.P. conducted the bioinformatic analyses. All authors analyzed the data. M.T.M. and M.C.J. wrote the manuscript with input from all authors. M.C.J. supervised the work.

## Competing interests

Princeton University, Stanford University, and the University of York have submitted a provisional patent application on aspects of the findings.

## Data and material availability

all data are available in the main text or the supplementary materials.

## Supplementary Materials

Materials and Methods

Figures S1-S6

Tables S1-S2

References (*1-45*)

