## Supplementary Materials for "Assembly of the algal CO_2_-fixing organelle, the pyrenoid, is guided by a Rubisco-binding motif"

**This PDF file includes:**

Materials and Methods
Figs. S1 to S6
References (*1-45*)

**Other Supplementary Materials for this manuscript include the following:**

Tables S1 to S2

### Materials and Methods

#### Strains and culture conditions.

The *Chlamydomonas reinhardtii* stain CC-4533 (*33*) was the wildtype for all experiments (hereafter WT) and parent for all genetic transformations. The fluorescently-tagged strain showing the native localization of SAGA1 was *saga1*-paro$^R$;SAGA1-Venus-3xFLAG-hyg$^R$ (*14*). All strains were maintained at room temperature (~22°C) under very low light (<10 µmol photons m$^{-2}$ s$^{-1}$), on solidified Tris-acetate-phosphate medium (TAP + 1.5% agar), pH 7.4, using a revised trace elements recipe for increased growth (*34*). Medium was supplemented with paromomycin at 2 µg mL$^{-1}$ for all strains (except WT), and additionally with 6.25 µg mL$^{-1}$ hygromycin for the SAGA1-Venus strain.

All experiments were conducted on photo-autotrophically grown cells. Liquid cultures were primed with a loopful of TAP-agar grown cells not older than 2 weeks resuspended into Tris-phosphate medium (as TAP above, but without acetate) to a starting concentration less than 10$^5$ cells mL$^{-1}$. Cultures were maintained in an orbital incubator-shaker (Infors) with controlled conditions: 130 rpm, continuous cool white fluorescent light at ~175 µmol photons m$^{-2}$ s$^{-1}$, 22°C, air enriched with 3% $CO_2$ v/v for faster growth and rescue of *saga1* and *epyc1*. Culture volume for Rubisco extractions was ~500 mL, for co-immunoprecipitations ~250 mL, for imaging and western blots ~50 mL. Cells were grown in conical flasks with a total capacity at least 4x that of the volume of the medium. Culturing time allowed for at least 6 rounds of mitotic division. Cell densities were not allowed to exceed 10$^7$ cells mL$^{-1}$ at any point in time and were sub-cultured accordingly. Cell densities were measured using a Countess II F automated cell counter (Thermo Fisher Scientific).

For most experiments, cells were acclimated to air-level $CO_2$ concentrations for 6 hours before harvesting, to maximize expression of the $CO_2$-concentrating mechanism and packaging of Rubisco into a pyrenoid (*35*, *36*). Cultures destined for confocal imaging were acclimated overnight (~16 hours). Acclimation to air-level $CO_2$ was performed by pelleting high-$CO_2$ grown cells (1,000 *g*, 10 min, RT), followed by gentle resuspension by agitation in fresh air-equilibrated TP medium, before transfer to an air-equilibrated chamber of the same orbital incubator-shaker (agitation, light, and temperature as above). $CO_2$ concentration was periodically monitored with a

$CO_2$ sensor (CO2Meter). All experiments aimed for a cell density at the time of harvesting of ~2-4 x$10^6$ cells $mL^{-1}$.

**Co-immunoprecipitation and mass spectrometry analysis.**

Native protein complexes were extracted according to the protocol described in Mackinder *et al.* (*18*), with minor modifications. Briefly, all protein extraction steps were performed at 4°C in a cold room, using only fresh algal material. After harvesting (1,000 *g*, 5 min, 4°C), cells were washed 1x in ice cold TP, re-pelleted, and suspended in a 1:1 (v/w) ratio of ice cold 2x immunoprecipitation (IP) buffer (400 mM sorbitol, 100 mM HEPES, 100 mM KOAc, 4 mM $Mg(OAc)_2.4H_2O$, 2 mM $CaCl_2$), containing a protease inhibitor cocktail (cOmplete, Roche), and phosphatase inhibitors (2 mM NaF, 0.6 mM $Na_3VO_4$). To ease the grinding, the cell slurry was transformed into frozen droplets of ~5 mm diameter by slowly releasing the cell/buffer mixture into liquid $N_2$ through a fine-tipped transfer pipette held ~15 cm above the cryogenic liquid. Releases were timed so as to avoid clumping of not fully frozen material. Each assay used ~1g of cell/buffer mixture. Mass spectrometry analysis was performed at the Stanford University Mass Spectrometry facility, as previously described (*18*). Raw spectral counts are given in Supplementary Table S1.

Minor deviations from (*18*) were: a 50/50 mixture of Dynabeads protein A and protein G was used; incubation was with anti-SAGA1 antibody (YenZym); protein complexes bound to magnetic beads were released by boiling for 1 minute; denatured protein samples were run on denaturing Tris/glycine gradient gel (4-15%), and stained with EZBlue (Thermo Fisher Scientific).

**Immunoblot analysis.**

Total proteins were extracted as follows. 10 mL cell suspensions were pelleted (3,500 *g*, 10 min, 4°C), resuspended in 300 µL lysis buffer (5 mM HEPES-KOH, pH 7.5, 100 mM dithiothreitol, 100 mM $Na_2CO_3$, 2% SDS, 12% sucrose, cOmplete protease inhibitor cocktail), transferred to a microcentrifuge tube, and heat denatured in a thermomixer (37°C, 10 min, 750 rpm). Lysate was clarified (16,000 *g*, 5 min, 4°C), aliquoted, flash frozen in liquid $N_2$, and stored at -80°C until analysis on SDS-PAGE.

Gel loading was normalized by total chlorophyll *a*+*b* content. Pigments were extracted from 50 µL cell lysate with 2 mL 100% methanol. Chlorophylls contained in the clarified extract

(16,000 $g$, 2 min) were quantified according to the following equations: chl. $a$ ($\mu g\ mL^{-1}$) = 16.29 $A_{665}$ - 8.54 $A_{652}$; chl. $b$ ($\mu g\ mL^{-1}$) = 30.66 $A_{652}$ - 13.58 $A_{665}$ (*37*), after correction for $A_{750}$. Absorbances were measured in a SmartSpec Plus spectrophotometer (Bio-Rad).

Proteins were separated by size on a denaturing Tris/glycine gradient gel (4-15%, Criterion TGX, Bio-Rad; 90V constant, 105min), transferred to 0.45µm PVDF membrane (Immobilion-P, MilliporeSigma) using a wet electroblotting system (Criterion Blotter, Bio-Rad) and Towbin buffer (20% methanol, 25 mM Tris, 192 mM glycine, 20% v/v methanol, 0.05% SDS), at 30V constant overnight.

For immunoblot analysis, membranes were blocked in TBS + 0.1% Tween-20 (TBST) containing 5% non-fat dry milk for 1h at RT or overnight at 4°C, under gentle agitation. Incubations with the primary antibodies were performed in TBST containing 2.5% milk for 1h at RT or overnight at 4°C. Membranes were washed in TBST (4x, 10min, rocking platform) before incubation with the secondary antibody for 1h at RT. Membranes were washed again in TBST (4x, 10min). Immunoreactive proteins were detected using enhanced chemiluminescence (WesternBright ECL, Advansta) followed by X-ray film processing (CL-Xposure Film, Thermo Fisher Scientific; SRX-101A, Konica-Minolta).

Primary antibodies were obtained from YenZym (anti-SAGA1 and anti-EPYC1) and MilliporeSigma (monoclonal anti-FLAG M2 antibody). The polyclonal anti-Rubisco antibody was a generous gift from Howard Griffiths, University of Cambridge, UK. Goat anti-mouse IgG (H+L) and goat anti-Rabbit IgG (H+L) were from Thermo Fisher Scientific. Dilutions: anti-FLAG 1:2,500 + secondary 1:10,000; anti-SAGA1 1:2,500 + secondary 1:10,000; anti-EPYC1 1:5,000 + secondary 1:10,000; anti-Rubisco: 1:10,000 + secondary 1:20,000.

**Rubisco purification and quantification.**

WT Rubisco was extracted as follows. 500 mL cell cultures were harvested (~4,000 $g$, 15 min, 4°C), resuspended in 1.5 mL lysis buffer (50 mM Bicine, pH 8.0, 10 mM $NaHCO_3$, 10 mM $MgCl_2$, and 1 mM DTT) containing a protease inhibitor cocktail, and transferred to an ice cold 50 mL Falcon (water/ice slush). Cells were sonicated on ice in 30 sec bursts followed by 30 sec pauses with a microprobe set at 60% amplitude (Q125 + CL-18 probe, Q Sonica), until no intact cells were left. Progress of the lysis was monitored with a light microscope (400x). Total soluble proteins were isolated by centrifugation (16,000 $g$, 30 min, 4°C), and 650 µL of the clarified lysate was loaded on top of a thin-wall ultracentrifugation tube (Ultra-Clear, Beckman Coulter) containing 12 mL of a 10-30% sucrose gradient prepared with the lysis buffer. Gradients were made the previous day with a gradient maker (Biocomp Instruments) and left to equilibrate at 4°C overnight. Gradients were run at 37,000 rpm for 20h in an ultracentrifuge (Optima XE-100 + SW 41 Ti rotor, Beckman Coulter). 750 µL fractions were collected either with a piston gradient fractionator (Biocomp Instruments) or manually by gravity. Fractions enriched in Rubisco were identified by running 10 µL aliquots in 2:1 Laemmli buffer on SDS-PAGE, followed by staining with EZBlue (same conditions as detailed in Immunoblot analysis, above). Fractions with the highest concentration of Rubisco (bands at 55 and 15 kDa for the Rubisco large and small subunits, respectively) were pooled, and buffer was exchanged by dialysis at 4°C overnight (Slide-A-Lyzer 20k MWCO, Thermo Fisher Scientific) using the same buffer as the one for the two Rubisco-peptide binding assays (see Surface Plasmon Resonance and Peptide Tiling Array, below). Rubisco was concentrated to ~2 mg $mL^{-1}$ on centrifugal filters (Amicon Ultra-4 100K, MilliporeSigma) before use. Rubisco concentration was determined by Bradford assay (Quick Start Bradford Dye Reagent + BSA Standard Set, BioRad).

**Binding of free synthetic peptides to immobilized Rubisco measured by Surface Plasmon Resonance (SPR).**

The SPR experiment was performed on a Biacore 3000 (GE Healthcare), at constant 25˚C, using the proprietary Biacore Control Software v.4.1, embedded application wizards *"Surface preparation"* and *"Binding analysis"*, and GE's immobilization kit, buffers and consumables (no deviation from the manufacturer's instructions). Optimal pH of 4.5 for amine coupling of purified Rubisco onto a CM5 sensor chip was identified with the aid of the *"pH scouting"* script. Variable amounts of Rubisco were immobilized in three independent assays (fresh Rubisco from independent extractions), spanning 2,000 to 6,000 resonance units (RUs) using the *"Aim for immobilized level"* script. All peptides (see Fig. 2B) were synthesized by GenScript, with purity ≥85% and nitrogen content validated by analysis on an organic elemental analyzer. Binding assays were run in PBS-P+ buffer (20 mM phosphate buffer, pH 7.4, 2.7 mM KCl, 137 mM NaCl, 0.05% v/v P20 surfactant). The same buffer was used for peptide solubilization and Rubisco dialysis (see Rubisco purification, above). Lyophilized peptides were solubilized to a stock concentration of 2.5 mM, aliquoted, flash frozen liquid $N_2$, and stored at -80°C until needed. Binding response was measured at a peptide concentration of 1 mM, during a 3 min injection into the sensor's flow cells at a flow rate of 40 µL $min^{-1}$. Dissociation was measured while injecting buffer only, at a rate of 40 µL $min^{-1}$ for 2.5 mins. The chip surface was regenerated by flowing buffer for 5 min at a rate of 40 µL $min^{-1}$. Return to baseline was observed for all peptides, except the one corresponding to RBMP2's fourth instance of the motif which remained partially insoluble even after addition of DMSO, and was therefore discarded from further analysis (see Fig. 3A). Binding responses were normalized to 1,000 RUs of immobilized Rubisco to allow comparison across independent repeats and plotted on a log10 scale. Peptides lacking the predicted Rubisco binding motif were used as negative controls: GYFAVDHRPNLAILQGELGTKSESMDVRI and SKPAVDLRFYLEIGMQNTA.

**Binding of free Rubisco to immobilized synthetic peptides measured by Peptide Tiling Array.** Four peptide arrays (30x20 spots each on 15x10 cm cellulose membranes, SPOT synthesis (*38*)) were ordered from the Koch Institute for Integrative Cancer Research at MIT, Biopolymers and Proteomics Laboratory (Cambridge, MA). The arrays were composed of 18-amino-acid peptides that tiled across the full-length of each of the six Rubisco binding protein sequences, with a step size of three amino acids. Each peptide was represented by a single spot, except for EPYC1, which included a non-randomized duplicate in positions 499 to 598 on membrane#1. All other locations of peptides were randomized. EPYC1 (100 spots, 2 repeats), CSP41A (142 spots), and RBMP1 (217 spots), were arrayed on membrane#1. RBMP2, SAGA1, and SAGA2, each required a separate membrane (558, 537, and 594 spots, respectively). Membranes were incubated with 750-2,000 µg purified Rubisco and probed by anti-Rubisco western blot, as described above. The peptide corresponding to the very C-terminus of each protein does not accurately represent the Rubisco-binding motif in this assay, as the peptides are linked to the cellulose via their C-termini, eliminating the carboxyl group which appears to be important for binding to Rubisco. Binding intensity was quantified in ImageJ (*39*) by measuring the integrated density of a circle of constant area centered on each blot dot, after background correction on an inverted image (rolling ball radius set to 25 pixels). Binding intensity was normalized to the binding of positive controls to allow comparison across membranes (TRSVLPANWRQELESLRN (*20*)).

**Fluorescent protein tagging and confocal microscopy.**

Open reading frames of the two chloroplast proteins *FDX1* and Cre12.g498550 were cloned by Gibson assembly into the vector pLM005-Venus (KX077945.1), as reported by Mackinder *et al.* (*13*). A 677 nt long GeneArt® DNA fragment (Thermo Fisher Scientific) and pLM005-Venus were double digested with EcoRI-HF and PflMI (New England Biolabs) and ligated by T4 DNA ligase (16°C). Constructs were validated by Sanger sequencing (759 nt PCR product spanning the double digested 643 nt insert) and transformed in WT Chlamydomonas, as described in (*13*), with two modifications: a 4-step pulse electroporator (NEPA21, NEPAGENE) was used and 50 µg of carrier DNA (MP Biochemicals) was added to each transformation.

*RBMP1*, *RBMP2*, and *SAGA2* were cloned using homologous recombination based on protocols optimized for Chlamydomonas (*28*).

Chlamydomonas transformants were selected on TAP-agar + 20 µg $mL^{-1}$ paromomycin, and high expressors were screened on a fluorescence laser scanner. Expression of full-length fusion proteins was validated by immunoblotting, using an anti-FLAG M2 antibody (Figs. S3F and S5G).

Images were captured with a laser-scanning microscope (TCS SP5, Leica) using a 100x objective with a 1.46 numerical aperture. Venus and chlorophyll were excited by argon lasers at 514 nm and 561 nm, respectively; emission was collected at [525-550] nm and [620-670] nm, respectively. Zoom-in (3X) acquisition settings were identical for all strains. 2D median plane cross-section were captured at 200 Hz. Pinhole was set at 1 airy unit. Venus was captured on hybrid detectors (HyD), chlorophyll autofluorescence on photomultiplier tubes (PTM). Picture montages were done on ImageJ (*39*).

**Bioinformatic search for motifs and motif enrichment analysis.**

We used a point system to identify motifs in the genome. A WR or WK dipeptide was needed to be considered a potential motif (obligatory condition but no point attributed). All motifs were relative to W ('zero' position). A basic residue (R or K) in position -8 to -6: +1 point (no additional point if multiple instances at those three positions). A proline (P) in position -3 or -2: +1 point. An aspartic acid (D) or an asparagine (N) in position -1: +1 point. An aliphatic residue (L, I, V or A) in position +4: +2 points. Finally, an acidic residue (D or E) or a COOH-terminus in position +5: +1 point. Proteins containing one or more motifs are listed in Supplementary Table S2. To test the statistical significance of the motif enrichment in pyrenoid proteins, we used the Mann-Whitney *U* test to evaluate the difference between the two distributions shown in Fig S5A, excluding the six proteins we originally noticed the motif in to avoid our observations biasing the result. The two distributions are different ($p = 0.047$).

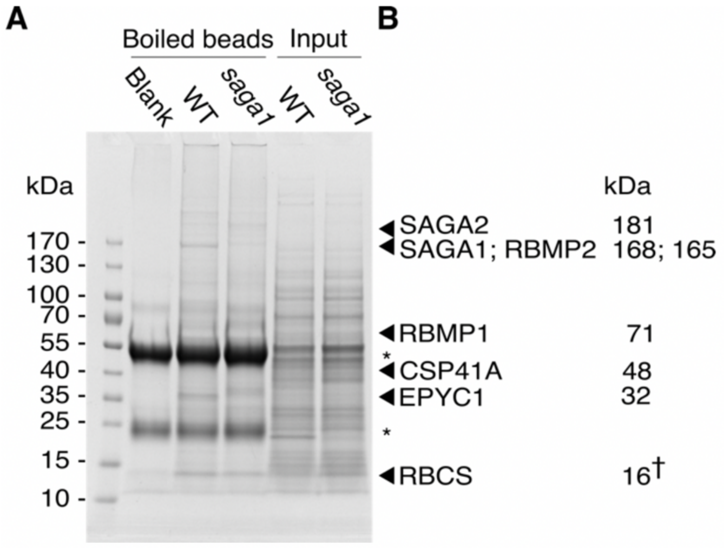


**Fig. S1. A polyclonal antibody raised against the pyrenoid protein SAGA1 interacts with at least five other pyrenoid proteins.**

(**A**) Coomassie-stained gel from the anti-SAGA1 immunoprecipitation experiment. First three lanes (identical to Fig. 1D): boiled beads alone (Blank), boiled beads after immunoprecipitation from wildtype (WT) and *saga1* lysates, respectively. The small and large chains of immunoglobulins used in the assay are labelled (*). Last two lanes: cell lysate input into immunoprecipitation experiment. (**B**) Calculated molecular weight (MW) of immunoprecipitated pyrenoid proteins, based on the full-length sequence (*40*), except for RBCS (†) for which the sequence of the mature protein is known. The MW was not adjusted for predicted chloroplast transit peptides.

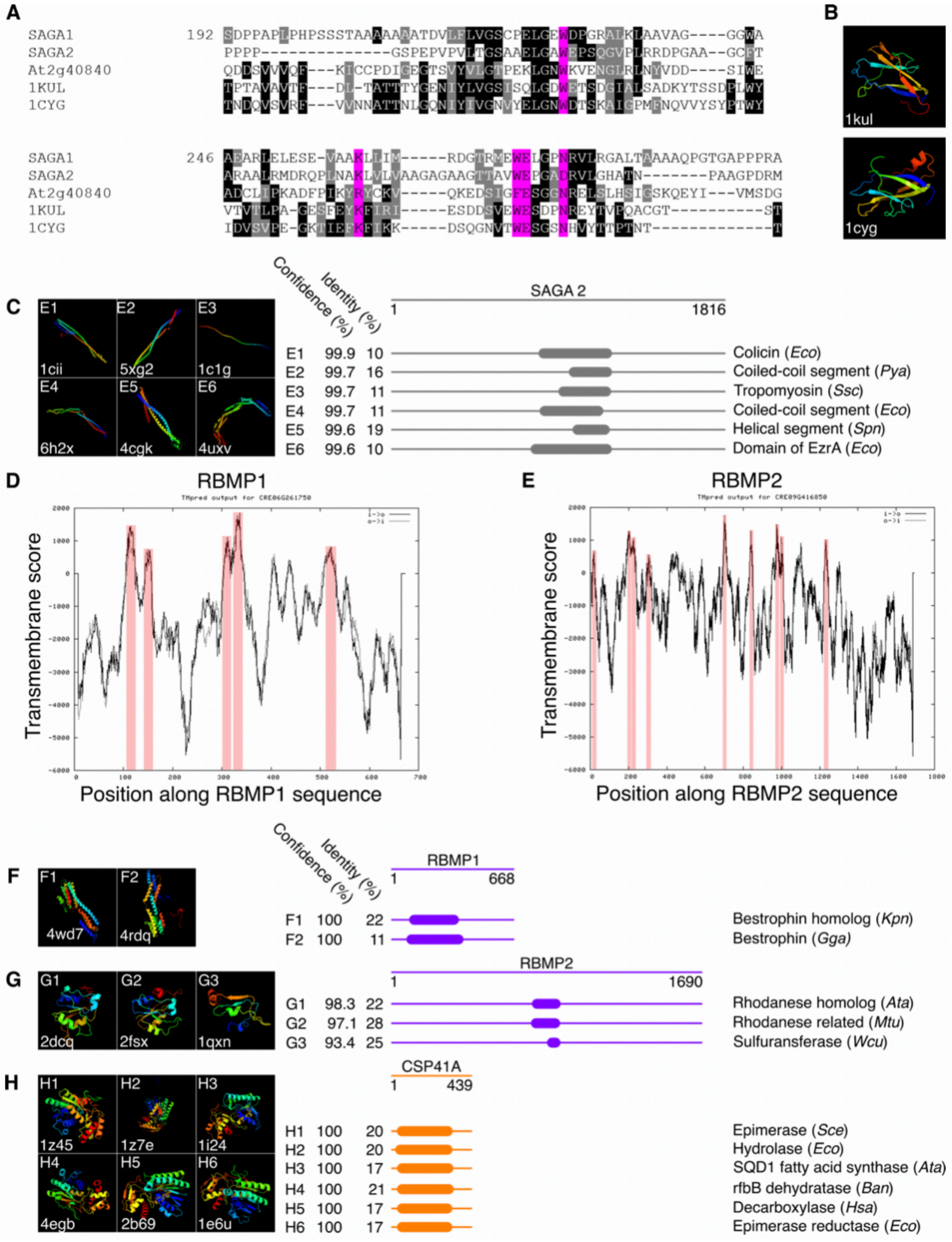
A
SAGA1
SAGA2
At2g40840
1KUL
1CYG
192
246
B
1kul
1cyg
C
Confidence (%)
Identity (%)
SAGA 2
1
1816
E1 99.9 10 Colicin (Eco)
E2 99.7 16 Coiled-coil segment (Pya)
E3 99.7 11 Tropomyosin (Ssc)
E4 99.7 11 Coiled-coil segment (Eco)
E5 99.6 19 Helical segment (Spn)
E6 99.6 10 Domain of EzrA (Eco)
1cii
5xg2
1c1g
6h2x
4cgk
4uxv
D
RBMP1
Transmembrane score
Position along RBMP1 sequence
E
RBMP2
Transmembrane score
Position along RBMP2 sequence
F
RBMP1
1
668
F1 100 22 Bestrophin homolog (Kpn)
F2 100 11 Bestrophin (Gga)
4wd7
4rdq
G
RBMP2
1
1690
G1 98.3 22 Rhodanese homolog (Ata)
G2 97.1 28 Rhodanese related (Mtu)
G3 93.4 25 Sulfuransferase (Wcu)
2dcq
2fsx
1qxn
H
CSP41A
1
439
H1 100 20 Epimerase (Sce)
H2 100 20 Hydrolase (Eco)
H3 100 17 SQD1 fatty acid synthase (Ata)
H4 100 21 rfbB dehydratase (Ban)
H5 100 17 Decarboxylase (Hsa)
H6 100 17 Epimerase reductase (Eco)
1z45
1z7e
1i24
4egb
2b69
1e6u

**Fig. S2. Functional predictions for pyrenoid proteins immunoprecipitated by the anti-SAGA1 antibody.** (**A**) SAGA1 and SAGA2 contain a predicted starch binding domain (belonging to the ubiquitous CBM20 family). Conserved residues predicted to be involved in starch recognition are highlighted (magenta). Numbering is for SAGA1. The partial alignment around the starch binding domain includes examples spanning the entire tree of life: a land plant (*Arabidopsis thaliana*, At2g40840), a fungus *(Aspergillus niger*, 1kul) and a Gram+ bacterium (*Geobacillus stearothemophilus*, 1cyg). (**B**) Crystal structures of two starch binding domains from (A). (**C**) Predicted regions of structural similarity between SAGA2 and experimentally determined protein structure data deposited in the Protein Data Bank (*41*). Six predicted structures are shown alongside PDB template ID, confidence and identity percentages. The position of a predicted functional domain is shown along the protein length of SAGA2 (to scale). Abbreviations: *Eco* (*Escherichia coli*), *Pya* (*Pyrococcus yayanosii*), *Ssc* (*Sus scrofa*), *Spn* (*Streptococcus pneumoniae*). (**D**) Predicted transmembrane domains in RBMP1 (*42*). Bestrophins are calcium-activated ion channels, that assemble as homo-tetramers or homo-pentamers (*28*, *43*). (**E**) Predicted transmembrane domains in RBMP2 (*42*). (**F** to **H**) Predicted regions of structural similarity between RBMP1 (F), RBMP2 (G), and CSP41A (H) with data deposited in PDB, as for (C), above. Abbreviations: *Kpn* (*Klebsiella pneumoniae*), *Gga* (*Gallus gallus*), *Ath* (*Arabidopsis thaliana*), *Mtu* (*Mycobacterium tuberculosis*), *Wsu* (*Wolinella succinogenes*), *Sce* (*Saccharomyces cerevisiae*), *Ban* (*Bacillus anthracis*), *Hsa* (*Homo sapiens*).

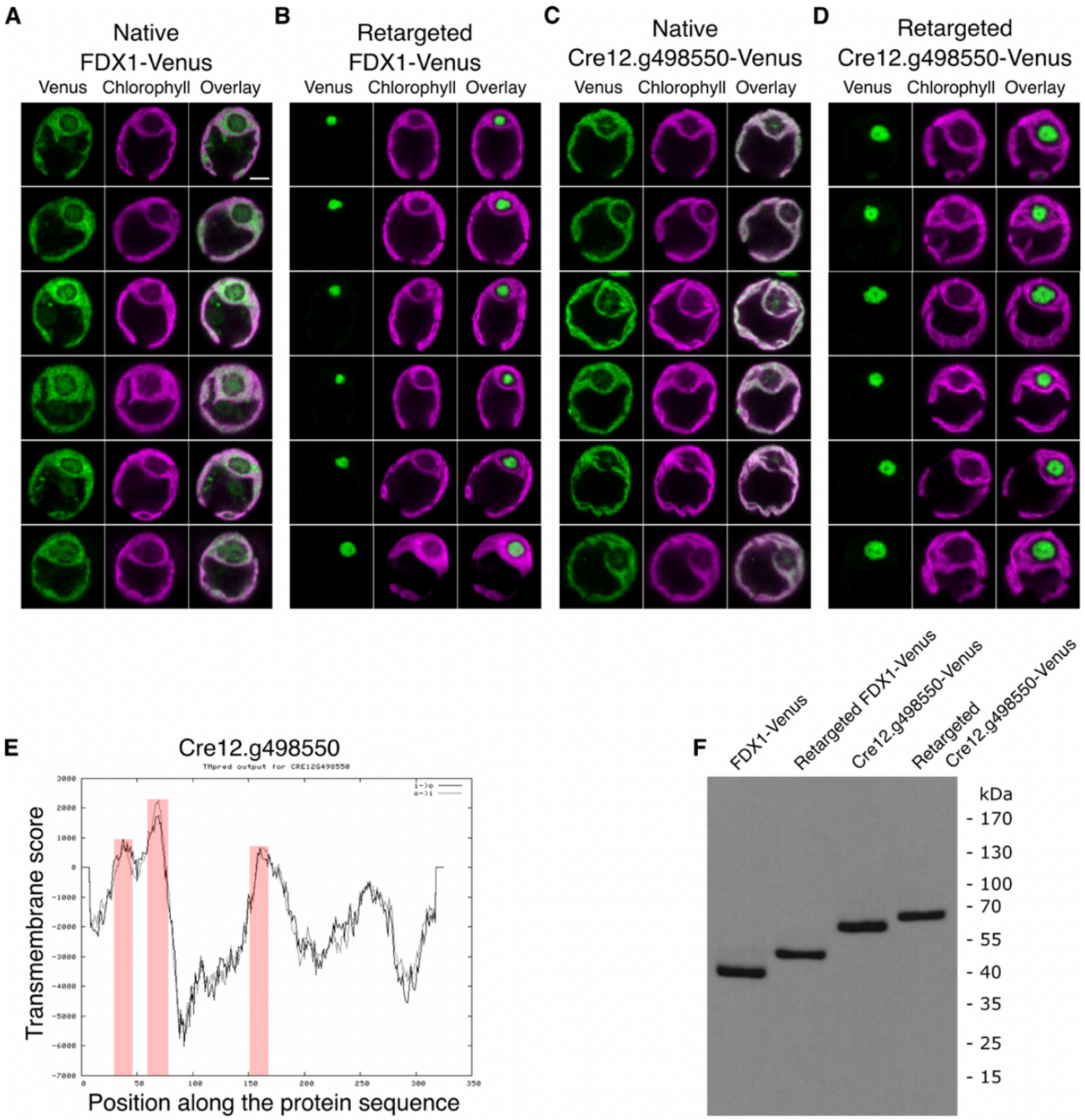


**Fig. S3. Subcellular localization of native and pyrenoid retargeted chloroplast proteins.** (**A**) Supplementary confocal images of the native localization of Venus-tagged ferredoxin 1 protein (FDX1). Scale bar, 2 μm. (**B**) Supplementary confocal images of FDX1-Venus with the C-terminal addition of three copies of the C-terminal SAGA2 motif. (**C**) Confocal images of the native localization of Venus-tagged Cre12.g498550, an uncharacterized chloroplast protein homologous to a Viridiplantae conserved magnesium protoporphyrin methyl-transferase involved in tetrapyrrole metabolism. (**D**) Confocal images of Cre12.g498550-Venus with the C-terminal addition of three copies of the C-terminal SAGA2 motif. (**E**) Predicted transmembrane domains of Cre12.g498550 (*42*). (**F**) Western blot validation of full-length expression of fluorescently tagged proteins. FDX1-Venus ≈13kDa; Cre12.g498550-Venus ≈35kDa; retargeting tag ≈8kDa.

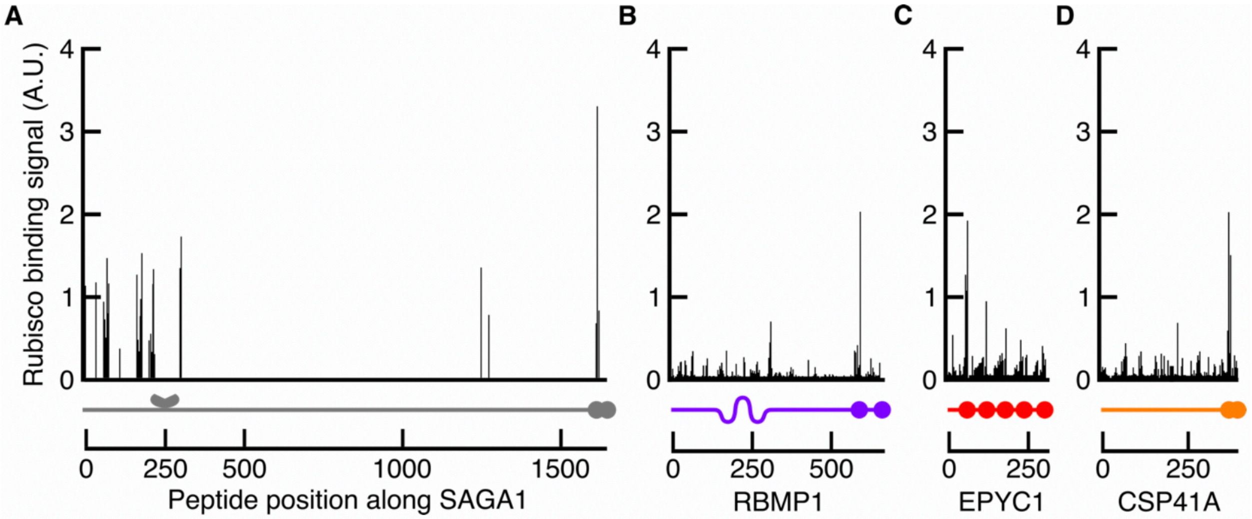


**Fig. S4. Rubisco-binding measured by peptide array**. (**A**) Array of 18 amino acid peptides tiling across the sequence of SAGA1, (**B**) RBMP1, (**C**) EPYC1, and (**D**) CSP41A. Arrays were synthesized and probed with Rubisco. Binding signal is normalized to a control EPYC1 peptide (same as for Fig. 3B and 3C, corresponding to one unit of binding). The positions of the predicted motifs are indicated to scale below each graph. The peptide corresponding to the C-terminus does not accurately represent the Rubisco-binding motif in this assay, as the peptides are linked to the cellulose via their C-termini, eliminating the carboxyl group which appears to be important for binding to Rubisco (based on the observation that internal instances of the motif are typically followed by an aspartic or glutamic acid, each of which carries a carboxyl group). The binding response was quantified by summing of all the pixel intensities in a constant circular region centered on each dot on the peptide array.

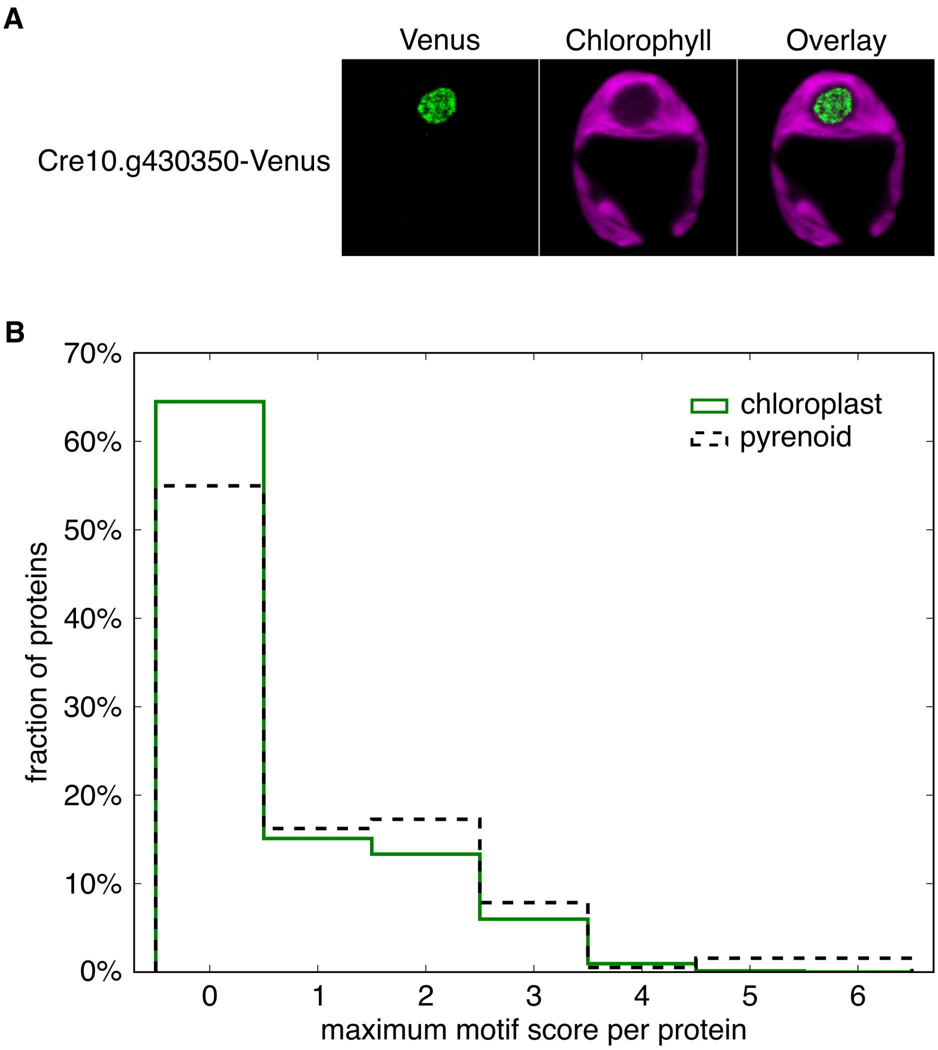


**Fig. S5. High motif scores were modestly enriched among pyrenoid proteome proteins relative to predicted chloroplast-targeted proteins. A.** Cre10.g430350, a motif-containing protein, localized to the pyrenoid matrix. **B.** The fraction of proteins with each motif score is shown. Pyrenoid: proteins found in the pyrenoid proteome (*17*). Chloroplast: proteins found in the chloroplast proteome (*44*) excluding proteins found in the pyrenoid proteome (see Table S2). The pyrenoid proteome has a distribution of motif scores that is shifted toward higher scores ($p = 0.047$, Mann-Whitney $U$ test; the six proteins used to define the motif [EPYC1, SAGA1, SAGA2, RBMP1, RBMP2, CSP41A] were excluded from the test to avoid biasing the result).

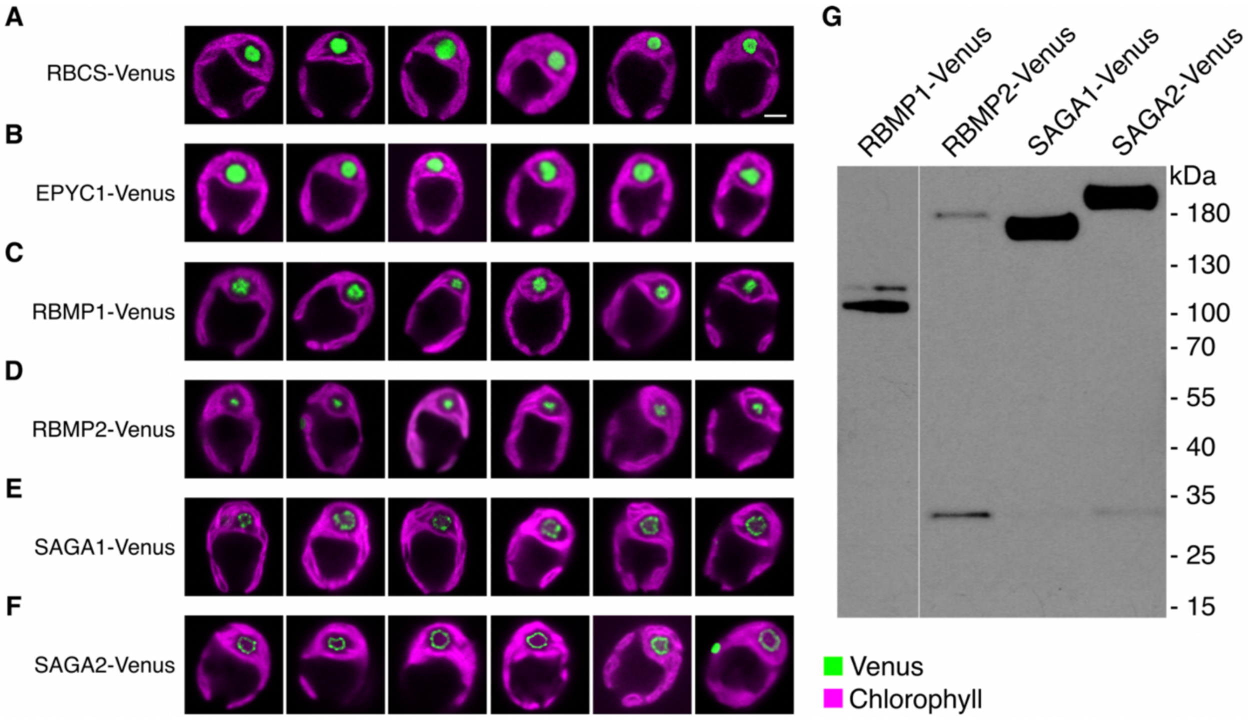


**Fig. S6. Subcellular localization of native pyrenoid proteins.** (**A**) to (**F**), Supplementary confocal images of the native localization of Venus-tagged RBMP1, RBMP2, SAGA1, SAGA2, EPYC1 and RBCS1, respectively. Chlorophyll autofluorescence delimits the chloroplast. Scale bar, 2 µm. (**G**) Western blot validation of full-length expression of fluorescently tagged proteins: RBMP1 ≈99kDa, RBMP2 ≈199kDa, SAGA1 ≈191kDa, SAGA2 ≈212kDa. For RBCS1-Venus and EPYC1-Venus, see (*18*).

**Table S1. (separate tab delimited txt file) Raw mass spectrometry counts of proteins coimmunoprecipitating with anti-SAGA1 antibody.** (See Fig. 1E) pyrenoid_proteome: + indicates presence in the pyrenoid proteome (*17*); - indicates absence. Rubisco_interaction_WD_score and Rubisco_interaction_Z_score are metrics of coimmunoprecipitation with RBCS-Venus-3xFLAG (see *18*).

**Table S2. (separate tab delimited txt file) *Chlamydomonas reinhardtii* proteins containing one or more instances of the motif.** See also Materials and Methods (Bioinformatic search for motifs). #motifs indicates the number of motifs in the protein. motif_scores indicates the motif scores in order from N to C terminus of the protein. motif_positions indicates the motif positions on the protein. motif_sequences indicates the sequence(s) identified as potential motif match(es). pyrenoid_proteome indicates presence or absence from the pyrenoid proteome (*17*); chloroplast_proteome indicates presence or absence from the chloroplast proteome (*44*). predalgo_protein_localization is the predicted protein localization based on the PredAlgo prediction algorithm (*45*).

20. S. He, H.-T. Chou, D. Matthies, T. Wunder, M. T. Meyer, N. Atkinson, A. Martinez-Sanchez, P. D. Jeffrey, S. A. Port, W. Patena, G. He, V. K. Chen, F. M. Hughson, A. J. McCormick, O. Mueller-Cajar, B. D. Engel, Z. Yu, M. C. Jonikas, The structural basis of Rubisco phase separation in the pyrenoid.

30. J. A. Raven, J. Beardall, P. Sánchez-Baracaldo, The possible evolution and future of $CO_2$-

concentrating mechanisms. *J. Exp. Bot.* **68**, 3701–3716 (2017).
